# Growth under constraints: root tip development controls trade-offs between speed and mechanical efficiency

**DOI:** 10.64898/2026.05.14.724970

**Authors:** Lionel X Dupuy, Jiaojiao Yao, Gloria de las Heras-Martinez

## Abstract

**Background and Aims:** Relationships between root tip development, particularly tip shape, are thought to play a key role in enabling plants to overcome soil mechanical resistance. However, correlations between root tip morphology and the ability to penetrate compact or hard soils have remained inconsistent.

**Methods:** This study applies a new quantitative framework to growth kinematics data from six plant species, analysing the trade-offs between frictional energy loss, growth stability, and reduced root elongation rates.

**Key Results:** The study shows a shorter root elongation zone can reduce the fraction of the mechanical energy lost to friction, but this is done at the expense of the elongation rate. A sharper tip or increased radius can help roots maintain the elongation rate at no energetic cost, but these strategies come with the cost of growth instability (tortuous roots) and decrease in specific root length respectively.

**Conclusions:** During establishment, root strategies may therefore occupy a 2-dimensional trait space in which the mechanical efficiency of growth is balanced against the explorative-exploitative trade-off.

## Introduction

The radius of a plant’s roots has often been linked to its ability to penetrate hard soils. This belief stems from the observation that radii tend to increase with soil mechanical resistance (Croser *et al*. 2000; Potocka and Szymanowska-Pułka 2018; Wang *et al*. 2021). Yet it remains unclear how a larger root diameter alone, if merely a matter of scaling, may improve the mechanical efficiency of growth. Any increase in growth forces from the tissue would be matched by those forces needed to deform a larger volume of soil. There is also the possibility that the radial expansion of root tissues occurs passively, as a result of increased axial forces, but computational models indicate the phenomenon may be insufficient to explain the variations in root diameters observed experimentally (Mimault *et al*. 2023).

Increases in root diameter may reflect deeper changes taking place during the development of the root apical meristem while growing in soil of varying resistance to penetration. Changes in root radius have notably been associated to modifications of the root anatomy (Zhang *et al*. 2024). Because variations in turgor pressure across root tissues are usually very modest, even in response to increases in soil mechanical resistance (Clark *et al*. 1996), the observed increases in root diameter and associated anatomical changes in harder soils may instead represent an adaptive response to enhance rigidity to reduce deflection. For example, reduced cortical area of maize roots has been linked to deeper rooting (Vanhees *et al*. 2020).

The reduction of the resistance to friction has been linked to an ensemble of other traits. The mucilage deposited at the interface with soil are believed to reduce the coefficient of friction of the root with the surrounding soil (Iijima *et al*. 2004; Rosskopf *et al*. 2022). Likewise, the root cap is known to facilitate the penetration of the root (Vollsnes *et al*. 2010), and it has been shown that border cells in the root cap contribute to this effect by detaching from the root cap when pushed through the soil (Bengough and McKenzie 1997). Root hairs help anchor the base of the root, enabling it to withstand the axial forces acting on the tip (Bengough *et al*. 2016). Movements such as twisting, circumnutation and tip deflections have also been linked to reductions in the resistance to penetration, through mechanisms that reduce axial friction or to allow growth through paths of least resistance in the soil (Mckenzie *et al*. 2013; Martins *et al*. 2019; Taylor *et al*. 2021; Leuther *et al*. 2025).

It remains unclear however, how the patterns of radial and longitudinal cell expansion at the root tip affects the generation of external forces applied upon the root. Recent research has highlighted the importance of root tip morphology in enabling plants to penetrate mechanically challenging soils. Colombi et al. (2017) found that sharper root tips elongated faster in compacted soil, but subsequent work carried out using root cap mutants showed that sharper root tips were less able to penetrate hard gel layers (Roue *et al*. 2020). These apparent contradictions stem from the fact that root mechanical efficiency of growth is a complex trait, and existing theoretical frameworks (Dexter 1987; Mckenzie *et al*. 2013) may fail to capture the trade-offs roots encounter while penetrating soil.

In this study, we propose to consider both friction and root deflections (hereafter referred to as growth instability) as the primary mechanical limitations to root growth (Figure 1) and introduced a theoretical framework that links both factors to root growth kinematics and, consequently, to the root elongation rate. We then used the theoretical framework to analyse growth strategies in response to physical constrains from the environment. Root growth kinematics data were obtained from published studies that reported changes to at least one physical properties of the growth substrate. These physical variables included some factors directly linked to growth biomechanics, such as water content and substrate strength, whereas others, such as temperature, were not directly related. The dataset was then complemented by new experiments conducted using agarose gel of varying penetration resistance. The analysis of the dataset revealed how roots may minimise the costs associated with slow growth, frictional resistance and growth instability.

**Figure 1.**
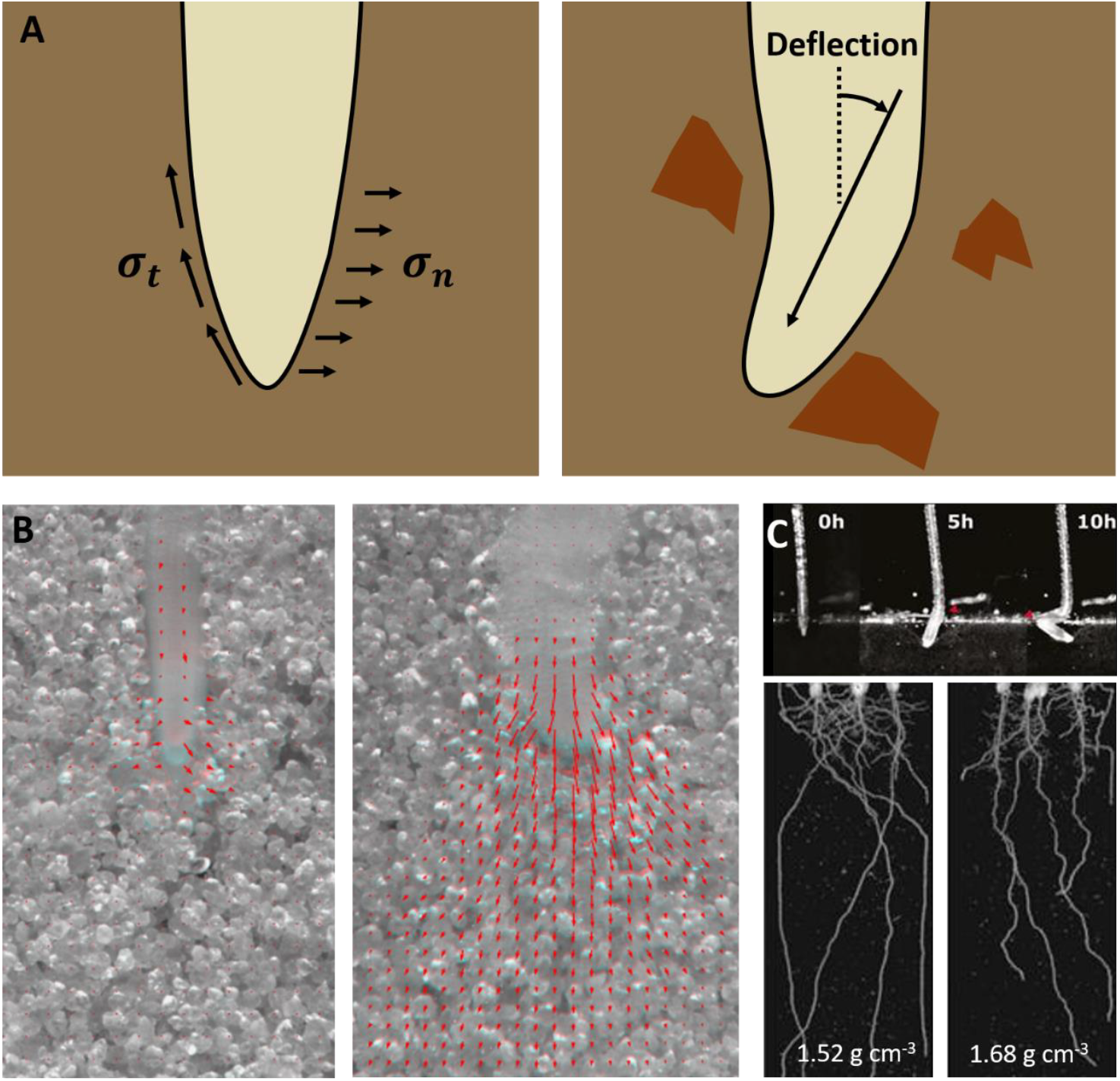
Illustration of the limitations to root growth resulting from the mechanical resistance from the soil. (A) Limitations are associated with the mechanical energy generated by the root to create a soil cavity (left). This energy is dissipated from the frictional stress *σ*_*t*_ generated by the soil slipping at the surface of the root and from the compression stress *σ*_*n*_ needed to expand the cavity. Another limitation is related to the ability of the root to maintain its growth trajectory (right), also referred here as growth stability. Growth instability leads to tortuous root trajectories, which reduce the efficiency of deep soil exploration. (B) Experimental data illustrating how increased soil friction leads to increase mechanical work from the root to form a cavity. The energy to deform soil is related to the deformation of the soil (red arrows), which was observed to be lower for maize plants with root caps (left) than when the root cap is absent (right). Data shared by Prof. Glyn Bengough from the published work of Vollsnes et al. (2010). (C) Illustration of growth limitations due to growth instabilities. When the axial forces resisting growth increases, roots are more prone to deflection because of buckling and bending (top). Image data reproduced from Roue et al. (2020). This results in growth trajectories that are less efficient at exploring the soil domain because of tortuosity (Bottom, soil bulk density indicated on the image). Reproduced from (Popova *et al*. 2016).

## Theoretical framework

### Root growth kinematics

The framework for describes energy dissipated by the root to form a cavity. The geometry of the root is assumed to be axisymmetric and embedded into the soil which is perfectly plastic and homogenous across the entire domain. Therefore, the model aims to represent an average mechanical behaviour of the root soil system. The framework relies on root growth kinematics, which offer a rigorous description of the motion of material points within a growing organ. The model represents the apical part of a root of length *L* [L]. That root is characterised by its elongation rate also equivalent to the tip velocity *V* [L T^−1^]. The cell expansion needed to produce this elongation rate occurs in a growth zone of length *L*_*e*_ so that *L*_*e*_ < *L*. The length *L* is assumed large enough so that its value does not affect any of the analyses and computations.

The root surface interacting with the soil is defined by a set of points **m** which coordinates are expressed as a function of the distance from the root tip *x* [L]. Hence, all quantities are defined in the moving reference frame of the root (Silk 1984): *x* = 0 indicate the position of the tip of the root and *x* = *L* is the base of the root fixed in soil,

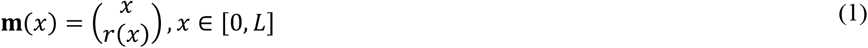

Here and throughout the remainder of this manuscript, vectors are represented using bold upright characters, while scalars are represented using italic characters. Next, we define the two-dimensional growth velocity, **v** = ∂_*t*_**m**. Using the chain rule, we have

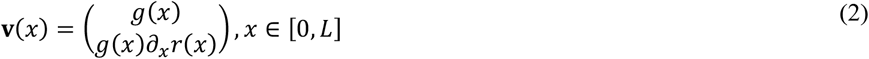

*g*(*x*) = ∂_*t*_*x* is a sigmoid like function describing displacement of cells relative to the root tip. It is typically obtained experimentally from image data (Beemster and Baskin 1998; Basu *et al*. 2007). It is such that beyond a distance *L*_*e*_ from the tip, it equals the velocity of the root tip *V* in the global reference frame *g*(*x*) = *V* for *x* > *L*_*e*_. [0, *L*_*e*_] is therefore the domain of the root where elongation occurs, whereas [0, *L*] is the entire root domain used for computations. The spatial derivative of *g* (∂_*x*_*g*), is the strain rate [T^−1^]. In this setting, the parameters of the functions *g* and *r* fully determine the root growth kinematics.

Next, we formulated the velocities of material points on the root envelop relative to the soil. The global root velocity **V** formulates the motion of material points on the root surface in the global reference frame and is used to compute the mechanical energy needed to form a cavity. The slip velocity 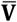 represents the motion of material points on the root envelop relative to the surrounding soil. It illustrates how a root reduces the energy lost by friction to the soil. **V** is obtained by correcting the growth velocity for the motion of the moving reference frame **v**,

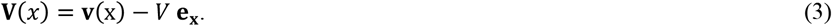

In the case of a rigid probe moving through soil, *g*(*x*) = 0 (the distance between any pair of material point in the probe is constant), and the velocity is constant and along the *x*-axis, **V**(x) = −*V* **e**_**x**_. In the case of a root tip, growth reduces the axial component and increases the lateral component of the global root velocity (Figure 2A). The slip velocity 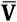 is obtained by calculating the difference between the velocity of the soil and that of the root along the tangential direction,

**Figure 2.**
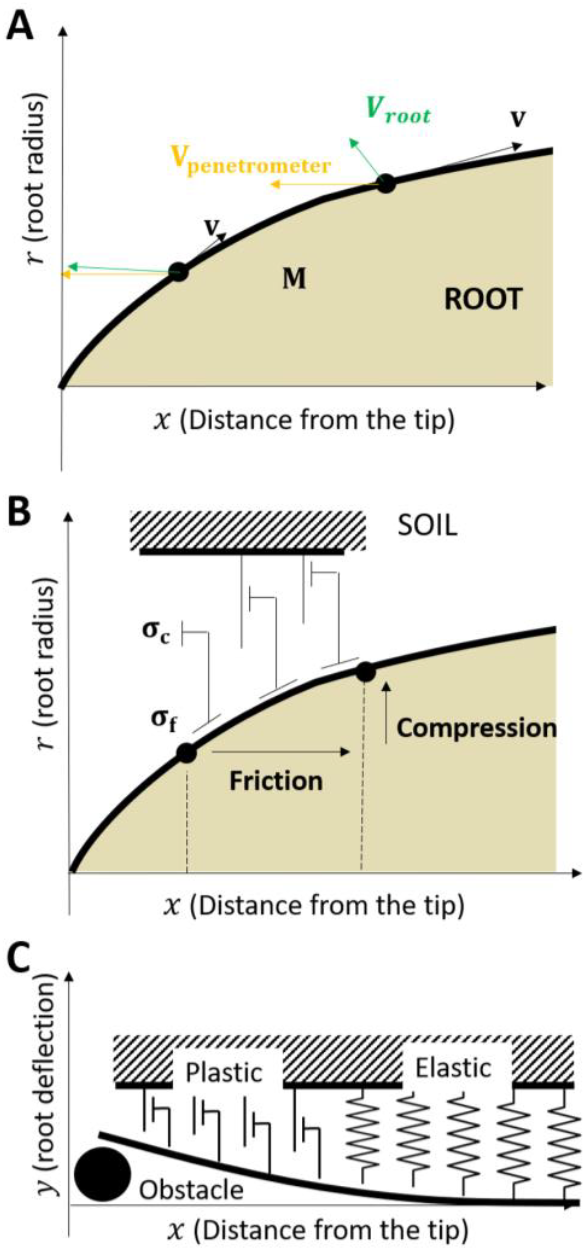
Mechanical interactions between soil and a growing root. (A) Growth kinematics describes the movement of material points *M* on the surface of the root through a velocity field. The problem is posed in the root reference frame so that each variable is defined spatially as a function of the distance to the root tip *x* and the distance from the root axis *y*. The velocity in the reference frame is denoted **v** (not capitalised), and is indicated in capital letter, **V** when expressed in the global reference frame. (B) The formation of the soil cavity requires the root to generate a pressure in the soil along the root, which at the interface with the root is represented through the normal stress *σ*_*n*_, and a stress tangential to the root surface *σ*_*t*_= *μ σ*_*n*_which is due to friction. Here *μ* is the friction coefficient. (C) The shape of root tips affects the ability of the root to bend through the second moment of inertia *I*. The deflection of the root can then be computer using beam on elastoplastic foundation.

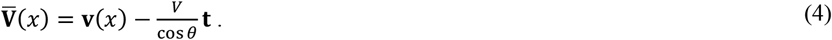

Whereas the slip velocity of a rigid probe moving through soil depends solely on its angle and speed, the slip velocity of a root progressively decreases with distance from the tip, thereby reducing the friction it experiences. The unit vector tangent to the envelop **t** is defined from the velocity vector **v**. The normal vector **n** is then defined as the unit vector perpendicular to the tangential vector and pointing outward of the root surface,

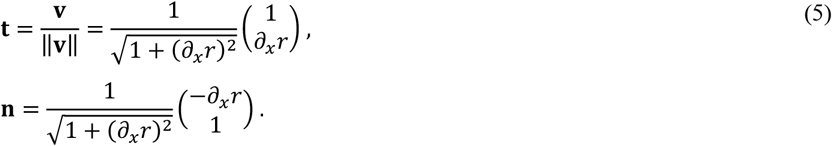

The magnitude of the velocity vector is defined as

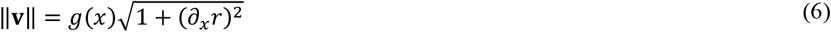

The angle *θ* of the root envelop with the *x*-axis is defined through trigonometric functions

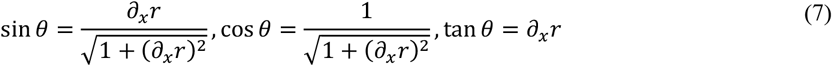

### Mechanical resistance to cavity formation

The displacement of material points along the root is resisted by the soil through two distinct mechanisms. First, the root must deform the soil to create a cavity. For this to occur, growth must apply a mechanical stress *σ*_*n*_ [M L^−1^ T^−2^] normal to the root surface, i.e. **σ**_**n**_ = *σ*_*n*_ **e**_**n**_. Second, the movement of root material points relative to the soil induces friction stress *σ*_*t*_ tangential [M L^−1^ T^−2^] to the root surface so that **σ**_**t**_ = −*σ*_*t*_ **e**_**t**_. We assume the normal yield stress needed to expand the cavity is constant, as determined for example from the cohesion of a plastic soil (Ruiz *et al*. 2015). The tangential stress follows the Coulomb model,

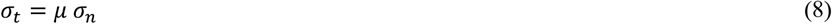

### Mechanical energy required to form the soil cavity

The resistance of the soil experienced by the root due to friction was formulated as the mechanical work (energy, [F L]) needed to deform the soil, because the quantity can more readily be linked to growth kinematics functions *r* and *g* in equations 1 and 2 (Figure 2B). For this, we consider the mechanical power (energy per unit of time, [F L T^−1^]) exerted by a slice of soil of unit length to move from position *x* to position *x* + *dx*,

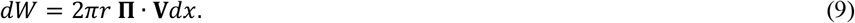

Here **Π** is the mechanical stress vector defining the resistance of the soil to root movement **Π** = (*μσ*_*n*_, *σ*_*n*_). To simplify the formulations, *σ*_*n*_ and *μ* are assumed to be independent of time and deformation of the soil. We used the convention that the energies and powers transferred to the soil are positive. Combining equations 3 and 9, we obtain

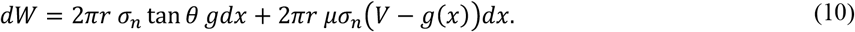

The mechanical power exerted by the soil is therefore the sum of a compression term and a friction term. The total mechanical power needed by the plant to produce a unit length of root is obtained by summing the contribution of all the root sections *dx* of the growth zone,

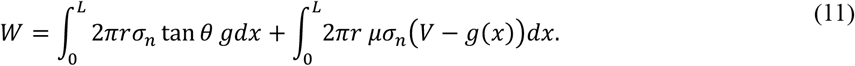

Since the mechanical power is the scalar product of force a velocity, we can therefore determine the force *F* [M L T^−2^] exerted by the root on the soil as

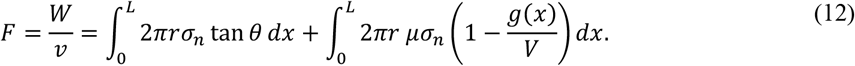

The first two terms of the right-hand side in equation 12 therefore represent the force for respectively the soil compression and the soil friction, and the term (1 − *g*/*v*) is the reduction of friction due to growth.

The change of variable *x* = *Rx*^′^ can be used to non dimentionalise the expression of the forces to express the mechanical resistance of the soil as a stress. The non-dimensional growth kinematic functions, defined on [0, *L*/*R*[ are such that *r*(*x*) = *Rr*′(*x*/*R*) and *g*(*x*) = *Vg*^′^(*x*/*R*). The force resisting root elongation is therefore,

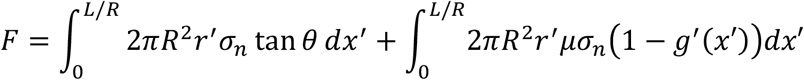

and model assumptions therefore result in the mechanical stress resisting penetration, *σ*_*p*_ [M L^−1^ T^−2^] being independent of the radius of the root,

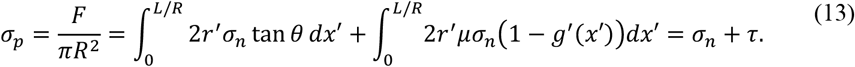

### Trait space for growth and form of the root tip

The next step is to define parameters that control the growth kinematic functions *r* and *g*. The growth kinematics function *g* is constructed assuming the strain rate is piecewise linear,

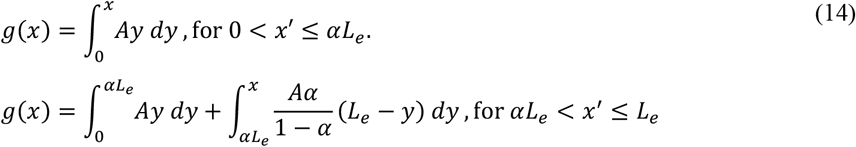

Two parameters define the strain rate. *A* is the growth acceleration parameter and *α* defines where the maximum strain rate is reached. The velocity of the root tip *V* = *αAL*_*e*_^2^/2. In the non-dimensional form,

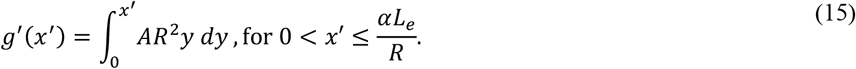

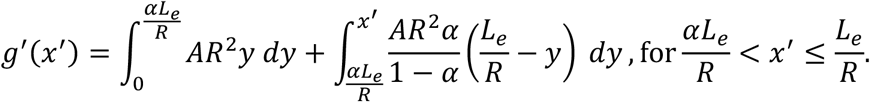

It is observed experimentally that the length of the growth zone *L*_*e*_ is the main variable responding to soil conditions such as water content (Sharp *et al*. 1988) or soil penetration resistance (Croser *et al*. 1999). Therefore we assumed the growth acceleration parameter (*A*) and the fraction of the growth zone where acceleration occurs (*α*) is constant for a given species. This hypothesis do not systematically hold for all plant responses, such as those induced by temperature.

The shape of the root tip *r* was defined using a truncated Hill function where the power, *γ*, determined the sharpness of the root tip. With the non-dimensional transformation *x* = *Rx*^′^ and *r* = *Rr*^′^,

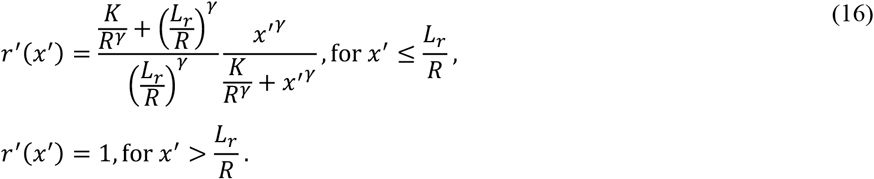

The formula allowed representation of both convex and concave tip shapes using the single parameter *γ*. In sensitivity analyses, the parameter *K* was set to the constant *L*/4 to maintain the shape of the first half of the root tip dominated by the power law *x*′^*γ*^. *L*_*r*_ was kept constant to *L*/2. All possible growth patterns and tip shapes affecting the resistance to root penetration could therefore be placed in a 2-dimensional parameter space comprised of the length of the elongation zone and the sharpness of the root tip (*L*_*e*_, *γ*).

### Growth stability

Here, we consider that the root tip is susceptible to deflection not only as a result of soil heterogeneity, but also because excessive axial forces or imperfections in root geometry can induce root tip deflection, a phenomenon we refer to as growth instability. To quantify growth instability of specific root tip geometries, we considered the magnitude of the deflection of the root tip in response to a lateral force of 1N. The radial displacement along the root is noted *y* and was such that the displacement at the base of the root is null *y*(*L*) = 0 and ∂_*x*_ *y*(*L*) = 0, and the displacement of the root at the tip, *y*(0), was used to quantify growth stability.

Within the framework of Euler-Bernoulli beam theory with elastic foundation the displacement of a linear structure such a root can be described by the Winkler foundation model (Teodoru and Muşat 2010). To account for plasticity in the deformation of the soil we solved the following equations (Figure 2C).

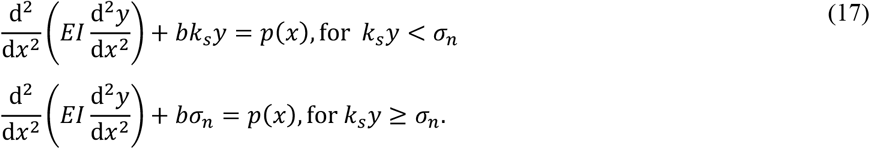

Here *k*_*s*_, is the modulus of subgrade reaction of the soil. *b* is the width of the soil bearing the root and *p* is the distribution of forces along the root.

## Materials and Methods

The model proposed in the previous sections represents a simplified description of the mechanical interactions that occur during root penetration of soil. Many factors may contribute to the forces experienced by the roots to deviate from model predictions. The normal forces assumed constant likely vary as a function of the root diameter and distance from the apex. Resistance to friction likely vary with tissue type, maturity water content. Time dependency of soil mechanical properties likely adds complex effects of growth kinematics on compression and friction stresses exerted on the root. Nevertheless, the model is a useful platform to interpret experimental variations. Experimental functions for *g* and *r* measure how much slipping is taking place along the root and likely correlate with energy losses to friction (equation 13). *r* alone likely correlates with the susceptibility of the root tip to deflect (equation 17). The following sections present the construction of a root growth kinematics dataset and the analyses employed to examine how variations in root growth kinematic parameters may relate to different root growth strategies.

### Experimental data on root growth kinematics

We compiled experimental data from the literature on root growth kinematics in response to mechanical constraints imposed by solid substrates. We could only find one study where tip shape and root growth kinematic of pea was measured in response of changes in soil strength (Croser *et al*. 1999; Kirby and Bengough 2002). We included data from Sharp et al. (1988) on the effect of soil water potential (maize roots), since reduction in the soil water potential reduce the root growth pressure and therefore links to biomechanical limitations. Finally, we incorporated two datasets examining root growth responses to temperature as control treatments, providing examples of growth responses that are not driven by mechanical limitations. These included the study of the effect of temperature on maize root tip growth from Pahlavanian and Silk (1988), and the study on the effect of temperature on *Arabodopsis thaliana* root tip growth from Yang et al. (2017).Data was extracted from figures extracted using WebPlotDigitizer (https://automeris.io/WebPlotDigitizer/).

Additional root growth kinematics experiments were also performed on seedlings of tomato (Solanum lycopersicum “Tres Cantos”, Fito, Spain), Cabbage (*Brassica oleracea* “Corazon de buey”, Fito, Spain), and wheat (*Triticum aestivum* rgt Tocayo, El Granero Integral, Spain). Roots were grown on agarose gel, since the substrate was shown to reproduce the penetration resistance of a broad range of soils (Clark *et al*. 1999). In addition, true *in situ* kinematic analyses have only measured soils deformations to date (Keyes *et al*. 2017). The agarose gel was obtained from powder (BP160-500 lot 220906, Fisher bioreagents, Spain) dissolved in deionized water at concentration of w/w, 1.5 %, 3.0 % and 4.5 % and subsequently heated at a temperature of 90°C until becoming transparent. The Agarose was then poured in the microcosm chambers (Liu *et al*. 2025) and cooled to room temperature. The resulting agarose gel had a mechanical strength of 74 kPa, 194 kPa and 323 kPa respectively (Yao *et al*. 2026). Before sowing into the microcosm chambers, seeds were first hydrated for one hour in sterilised water and then sterilised for 15 minutes in a 1% calcium hypoclorite solution (042548.30, Thermo Scientific, Spain). After numerous washes, seeds were placed in the microcosm chamber and then placed in an incubator at 23 ºC during 24h.

To obtain growth kinematics data, images of the root tip of seedlings were acquired every hour. The imaging system consisted of a BSI Prime Express CMOS camera (Teledyn, UK) collecting 16-bit images formed by 5X and 2X Mitutoyo Plan Apochromat Objective (Figure S1 A). Light was transmitted through the sample using a 3.2” TFT LCD panel, (1743 Adafruit, Mouser Electronics, Spain). The LCD panel was turned on only during image acquisition. Time-lapse images were acquired every 2 min over 1 h (30 time points, Figure S1 B). For each plant species and agarose concentration (1.5%, 3.0% and 4.5%), at least four biological replicates were obtained. We used the Kineroot software (Basu *et al*. 2007) to fit the root radius function *r*(*x*) and growth velocity function *g*(*x*) (Figure S1 C&D).

### Analysis of root growth kinematic data

Data was non-dimensionalised to enable the comparison of traits from species of different radii. To understand variations in the frictional resistance to growth, distances were normalised by the length of the root elongation zone *L*_*e*_. After applying the transformation *x*′ = *x*/*L*_*e*_, the normalised growth velocity function *g*′(*x*′) from all plant species collapsed onto a single curve (Figure S2). Then variations in the radius function becomes indicative of the surface of the root exposed to friction, since sharper root tips functions *r*’(*x*’) have smaller radii for a given slip velocity. To analyse variations in growth velocity, we normalised distances by the radius of the root *R*. Since the growth velocity scales with the square of the length of the elongation zone (*V* = *αAL*_*e*_^2^/2, equation 14), velocity was normalised by the square of the radius, *V*’ = *V*/*R*^2. Following this transformation, variations in the root elongation rate can be attributed to differences in both the length of the elongation zone and the growth acceleration parameter *A*. Growth stability was quantified as the lateral deflection of the tip to a constant force of 1N. The scaling of the lateral displacement *y* with the radius of the root is not straightforward. For example, in the case of a beam on an elastic foundation, deflections scale with the radius at the power 7/4 (Hetényi 1946). Therefore, normalisation was performed computationally. The root radius functions were therefore normalised to 1 mm diameter before computations of the lateral deflection of the root tip.

Root growth strategies were characterised for three traits (Γ_*i*_)_*i*≤3_ such that larger trait values are beneficial and smaller values are disadvantageous. The first of these traits is the growth velocity Γ_1_ = *V*, the second is the friction trait, Γ_2_ = 1/*τ*, and Γ_3_ = 1/*y*_*a*_ is the growth stability trait. To infer variations in the underlying growth strategy of the different plant species studied, we computed root traits’ multiplicative contrasts **κ** such that *κ*_*i*_ ∝ Γ_*i*_/(Γ_*j*_Γ_*k*_)_*j*≠*k*≠*i*_. Because each trait is normalised by the values of the other two, the resulting contrast emphasises the trait that dominates the others.

Min-max scaling was applied to the dataset of contrasts so that 0 ≤ *κ*_*i*_ ≤ 1 for all root tip in the dataset. To collapse the entire dataset of the study into a single plot, we identified 3 extreme species and conditions, each maximising one of the multiplicative contrasts.

This lead us to define 3 sets of multiplicative contrast 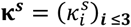, where by definition 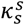 is the largest s^th^ multiplicative contrast of the entire dataset and 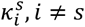, *i* ≠ *s* are the other two contrasts for that phenotype. For example, 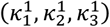 are the multiplicative contrasts of the phenotype that maximise velocity. A probability of similarity to the *s*^th^ extreme phenotype was then determined using a Softmax function,

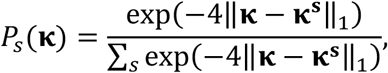

where ‖ ‖_1_ denotes the Manhattan norm. The probabilities *P*_*s*_ for each experiment in the dataset were then displayed on a single ternary plot.

One final question is what growth strategies plant species adopt to cope with increasing soil strength. To address this question, we first hypothesised that the cost associated with a given growth strategy is the sum of penalties for slow growth, frictional resistance, and deflection of the root tip, *E*_**β**_ = ∑ *β*_*i*_/Γ_*i*_. Because the error function is a linear combination of individual cost functions, simple formulas can be derived to infer the weight *β*_*i*_ a plant species allocate to each penalty (Supplementary Information). This approach also enables us to determine how the growth parameters (*L*_*e*_, *γ*) must change in order to maintain the same weight parameters *β*_*i*_ when the mechanical resistance of the soil increases (Supplementary Information). We therefore fitted the growth functions (equations 15 and 16) to determine the parameters *L*_*e*_, *γ* for each species and experimental conditions. The weights *β*_*i*_ were determined for roots grown in mechanically unconstrained conditions. The changes in growth parameters (Δ*γ*, Δ*L*_*e*_) were determined both from the experimental data and as predicted from equation 5 in the supplementary information. Since growth conditions are not always comparable, we compared the observed and predicted ratio Δ*γ*/Δ*L*_*e*_.

## Results

### Effects of root growth kinematics on the energy lost from friction

The newly developed mathematical framework couples root growth kinematics with mechanical analyses to estimate the energy dissipated by the root as it deforms the surrounding soil. Since the energy lost to friction depends on the root-soil slip velocity, it is possible to determine how growth kinematic enables to reduce frictional resistance to growth. Hence, a central result of this work is the equation 12 that quantifies this relationship. The assumptions made to obtain these equations are simpler than more computationally intensive methods to determine soil deformation around growing roots (Ruiz *et al*. 2017; Koren *et al*. 2024), but instead exploits cone penetrometer test formalism commonly used in soil sciences. It is easily verified that in the case where the object penetrating the soil is a conical probe of semi angle *α* (*θ* = *α* = cte in equation 12) and in the absence of growth (*g* = 0 in equation 12), the model predicts penetration resistance found in the literature (Mckenzie *et al*. 2013),

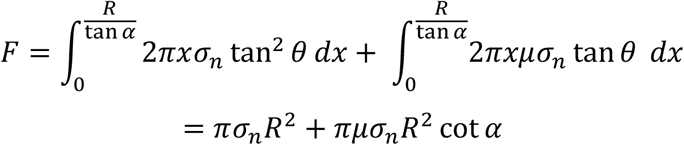

The mathematical framework also informs on factors that affect the mechanical efficiency of root growth. Beyond a certain distance from the root tip (*x*^′^ > *L*_*e*_/*R*) both the angle of the root surface (*θ*) and the slip velocity (1 − *g*^′^) vanish. One consequence of the model (as formulated in equation 13) is that the energy dissipated (*σ*_*p*_) is independent of the root radius *R*. The contribution of the compression of the soil always integrates exactly to the yield stress of the soil *σ*_*n*_. Although this property may not hold in natural soils or even in artificial growth substrate, compression stress most likely increases with root diameter because of the increased density of the soil around the root (Gregory *et al*. 2006). It therefore indicates that increases in root radius are likely associated with increased compression and friction costs.

Another property of the model is that the energy used to compress the soil is independent of the growth kinematics, and there is no mechanical energy gained or lost through changes in root radius. Since the yield stress *σ*_*n*_ is chosen independent on the strain rate (pure plasticity), the friction coefficient *μ* was independent of slip velocity, and *g*^′^ was likewise independent of tip velocity. As a consequence of these model properties, the theory predicts the energy required to create a unit length of root cavity is independent of the root elongation rate. Only the location where cell expansion occurs, the function *g*^′^, affected the frictional resistance to root penetration. Here too, the property may not hold for certain soils. For example, dependence on the root elongation rate may arise, for example if the time scale of the consolidation process matches that of the growth rate of the root. Viscous friction may control the shearing stresses at root tips where mucilage is present. These factors would introduce a time dependency on either or both the compression and friction energy, but are not considered in this theoretical framework.

Independently of model simplifications, the analysis told us that an important benefit of increasing root radius is the increase in the root elongation rate. The non-dimensional forms of the growth kinematic functions *r*^′^ and *g*^′^ in equations 15 and 16 show that roots with equal *γ* and *L*_*e*_/*R* loose equal frictional energy to the soil. Amongst this family of growth kinematic functions however, those with larger root radius (*R*) have a larger root growth zone (*L*_*e*_) and this results in an increased elongation rate since *V* = *αAL*_*e*_/2. Collectively, these results show only two parameters are needed to explore the trait space, the length of the growth zone and the sharpness of the root tip (*L*_*e*_ and *γ* respectively). These parameters generate a trait space with minimal dimension (Figure 3 A&B) which adequately represent the strategies a plant may adopt to overcome soil mechanical resistance. A smaller (larger) *L*_*e*_ value corresponds to slower (faster) root growth. Likewise, smaller (larger) *γ* values indicate that radial expansion occurs earlier (later) relative to longitudinal growth. Of these two parameters, only *γ* affects growth stability in equation 17, and simulations showed the effect is stronger than changes in root mechanical properties (Figure 3 C&D).

**Figure 3:**
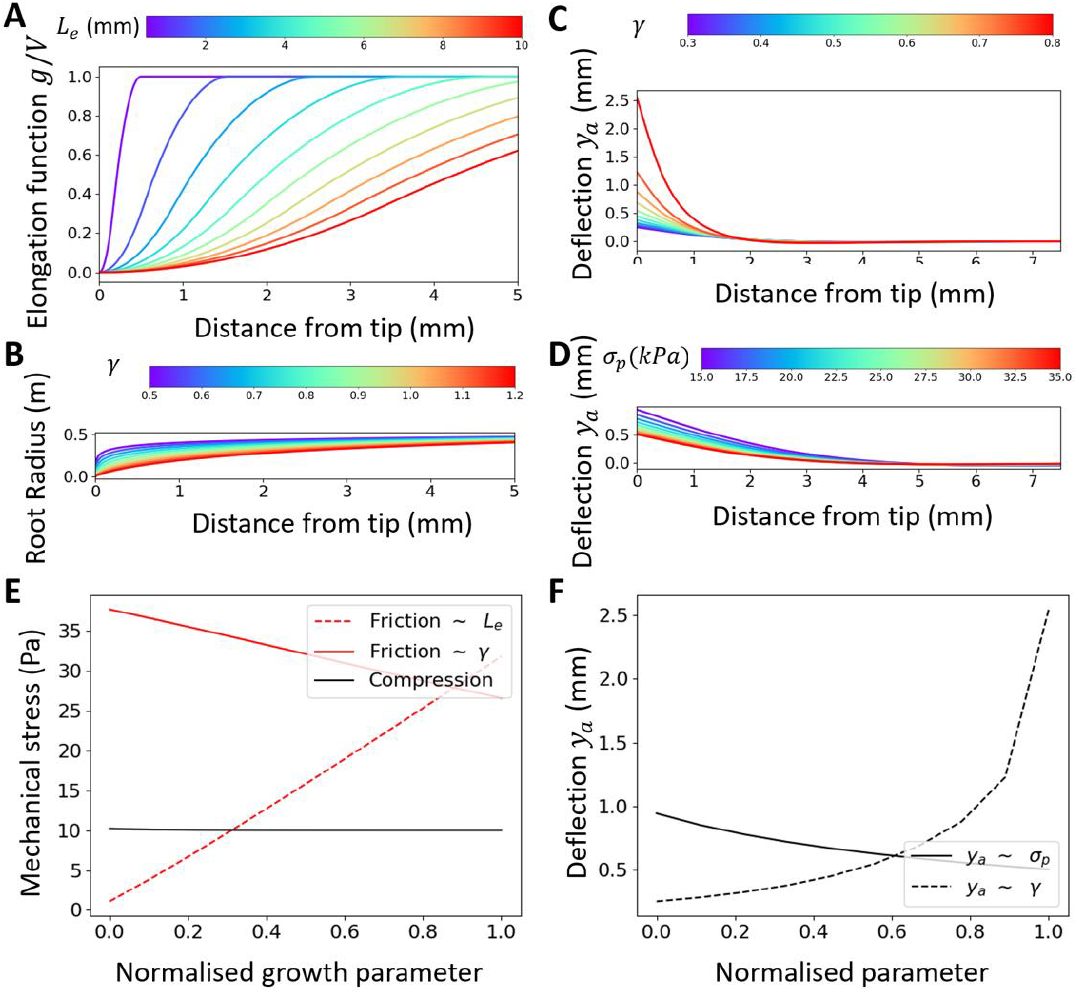
Root tip development and mechanical efficiency of growth. (A) Effect of the length of the growth zone *L*_*e*_ on the normalised velocity profile *g*^′^ = *g*/*v*. (B) Effect of the sharpness of the tip *γ* on the shape of the root tip. (C) effect of the sharpness of the root tip *γ* on the bending of the root. (D) Effect of the normal yield stress (*σ*_*p*_) on the bending of the root. The colour bars indicate the range of values for *L*_*e*_, *γ* and *σ*_*p*_ used for generating the plots. (E) compression (black line) and friction (red lines) stress fractions of the mechanical resistance opposing the growth of the root. Increasing the length of the growth zone *L*_*e*_ significantly increases the friction, while increasing the sharpness of the root tip decreases the friction experienced by the root. (F) Effect of the soil’s yield stress in compression *σ*_*p*_ (plain line) and sharpness of the root tip *γ* (dashed line) on the deflection of the root.

### Trade-off between reducing friction and maintaining a desirable growth trajectory

The reduction of friction due to growth kinematics, the factor (1 − *g*/*V*) in equation 12, can take values between 0 and 1. This indicates that frictionless growth can theoretically exists if radial expansion is completely separated from radial expansion. However, it is helpful to examine the trade-offs that confine root development within specific morphogenetic boundaries. The frictional resistance opposing root elongation depends on the sharpness of the root tip. Sharper tips, although subject to the same slip velocity as blunt ones, experience friction over a reduced surface area, which allows a reduction of the overall frictional forces resisting growth (Figure 3 E, red plain line). Reducing the length of the growth zone has a similar effect because shorter elongation zones concentrate friction at the apex, where root cross sections are the smallest (Figure 3 E, red dashed line). However, each of these responses comes with a trade-off. Increased sharpness result in the root being more susceptible to bending and multiple deflections (Figure 3 F). Shorter elongation zones on the other hand reduce the root elongation rate.

To analyse how plants manage these trade-offs, it is useful to consider which costs are minimised when particular growth kinematic parameters are observed. This was achieved by assigning penalties for reduced elongation rate (*β*_1_), mechanical friction (*β*_2_), and deviations in root growth trajectory (*β*_3_), and defining the overall error as the sum of these penalties. The relative weight given by a plant to each of these penalties was obtained directly from growth kinematic functions *r* and *g* fitted on experimental data. The resulting error functions (Figure 4) expressed the different trade-offs through the fitted penalty coefficient. Excessive values of the parameter *γ* create root tips that are very sharp. These configurations minimise energy lost to friction, but the resulting phenotypes are either inherently unstable—effectively pushing a soft, needle-like structure into a mechanically resisting medium (top left)—or extremely slow when *L*_*e*_approaches zero (top right). Likewise, low values of *γ* create root tips that are blunt and therefore mechanically stable, but when the length of the growth zone *L*_*e*_ is large the energy lost by friction is excessive (bottom right) and when *L*_*e*_ is small, the elongation rate is excessively small (bottom left).

**Figure 4:**
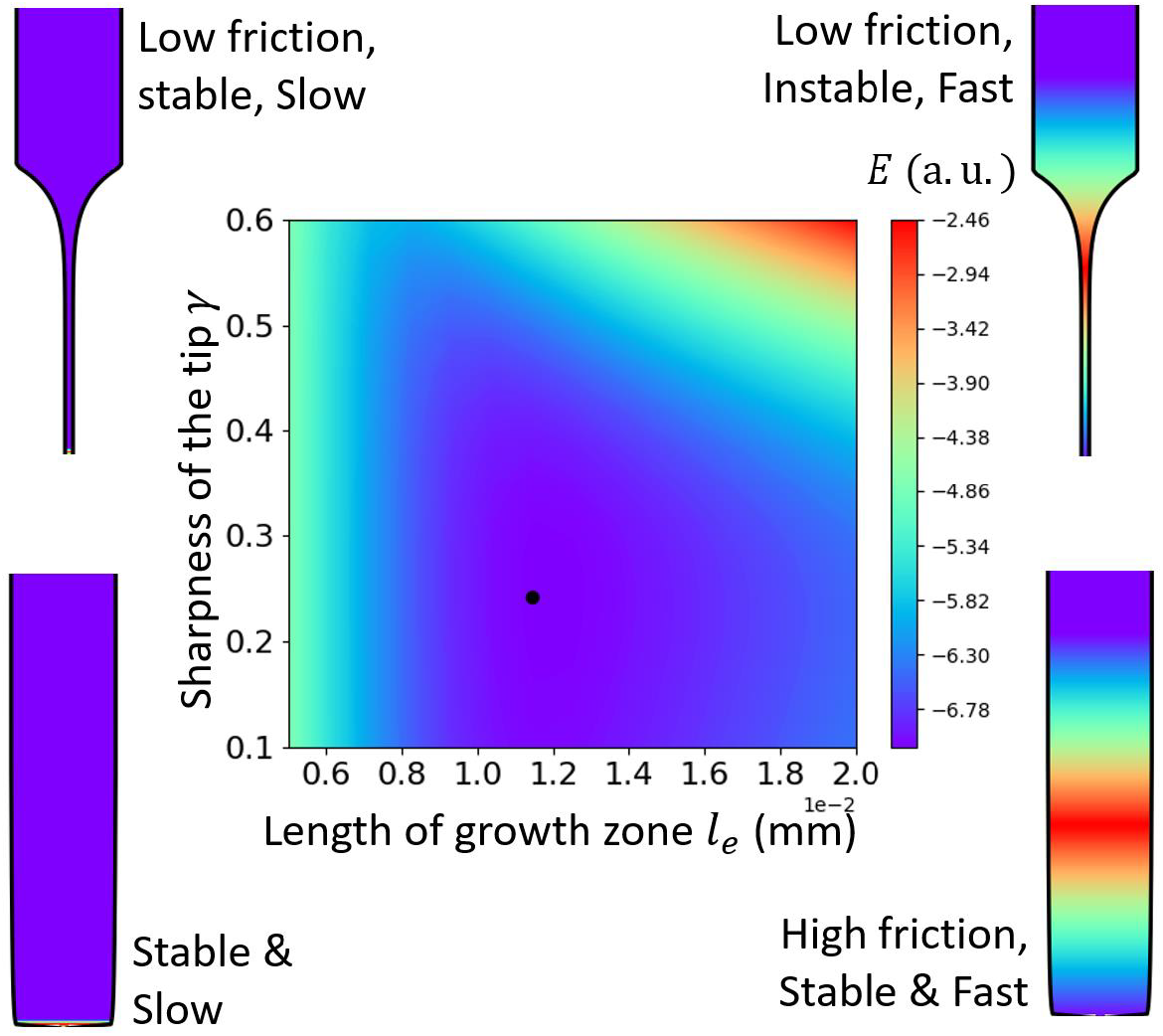
Cost function illustrating the trade-off between speed, stability and friction (centre). The top corners represent roots with the sharpest possible tips. These geometries cancel frictions but are slow (top left) or unstable (top right). The bottom corners represent completely blunt root tips. These geometries are either slow (bottom left) or high friction (bottom right). An optimal tip shape are roots which minimise a cost function *E* (here defined as in supplementary method) and witter with *L*_*e*_ ∈ [0.5 mm, 10 mm], *γ* [0.5,1.2] and *σ*_*p*_ ∈ [5 kPa,35 kPa].

### Variations in root growth properties

Root growth kinematics were subsequently determined for seven plant species: five crops (cabbage, wheat, tomato, pea, and maize), and the weed and model plant *Arabidopsis thaliana*. Experimental data were extracted from existing publication based on the availability of measured root radius and growth velocity profiles under at least two contrasting environmental conditions, including temperature, substrate water potential, and substrate mechanical strength.

The collected data revealed large variations in root growth kinematic parameters (Table 1). Many of these parameters scaled proportionally with root radius over approximately one order of magnitude. This included the growth acceleration parameter *A* (Pea 0.04 mm^-1^ h^-1^ and Arabidopsis 0.69 mm^-1^ h^-1^), the length of the growth zone *L*_*e*_ (Arabodopsis 1.22 mm and Maize 12.79 mm) or the half-saturation constant for the radius function *K* (Wheat 0.23 mm^*γ*^and Pea 2.46 mm^*γ*^). The growth acceleration zone parameter *α* was more conserved and less correlated to root radius (Tomato 0.23 and Cabbage 0.59). The tip shape parameter *γ* was related to the root radius but the magnitude of the variations was limited (Pea 0.30 and wheat 1.15). Parameters computed for an equivalent root diameter of 1 mm showed variation of about one order of magnitude, with Arabidopsis exhibiting the faster elongation rate (12.60 mm h^-1^), while wheat roots experience the lowest friction (3.13 kPa). The root tips of Arabidospsis plants deflected the least (0.09 mm) while those of wheat exhibited the largest deflections (65.45 mm). Such large variations arose because deflection scales with the second moment of inertia of the root cross-section, which itself scales with the fourth power of the root radius, resulting in variations spanning multiple orders of magnitude.

**Table 1:** Variation of root growth kinematics across a selection of plant species and in different environments. T is for temperature, C is for agarose concentration, ψ is for water potential, *σ* is for penetration resistance. Velocity, friction and deflections values are computed for a root of 1 mm in diameter.

| Species and treatment | $A$<br>mm<sup>-1</sup> h<sup>-1</sup> | $Le$<br>mm | $\alpha$ | $R$<br>mm | $K$ | $Lr$<br>mm | $\gamma$ | Velocity<br>$V$<br>mm h<sup>-1</sup> | Friction $\tau$<br>kPa | Deflection $y_a$<br>mm |
| --- | --- | --- | --- | --- | --- | --- | --- | --- | --- | --- |
| Arabidopsis (T = 15.0 °C) | 0.25 | 1.97 | 0.51 | 0.16 | 0.39 | 0.40 | 0.56 | 9.30 | 42.84 | 0.39 |
| Arabidopsis (T = 30.0 °C) | 0.69 | 1.22 | 0.35 | 0.12 | 0.24 | 0.28 | 0.67 | 12.60 | 30.42 | 0.09 |
| Cabbage ($\sigma$ = 74 kPa) | 0.10 | 3.77 | 0.59 | 0.41 | 0.38 | 1.67 | 0.84 | 2.41 | 25.67 | 6.31 |
| Cabbage ($\sigma$ = 194 kPa) | 0.10 | 3.23 | 0.55 | 0.48 | 0.65 | 3.98 | 0.67 | 1.29 | 11.77 | 20.67 |
| Cabbage ($\sigma$ = 323 kPa) | 0.09 | 3.23 | 0.41 | 0.48 | 0.65 | 3.44 | 0.77 | 0.81 | 9.43 | 26.86 |
| Maize (T = 19.0 °C) | 0.05 | 12.70 | 0.39 | 1.09 | 1.27 | 6.80 | 0.46 | 1.37 | 30.11 | 4.90 |
| Maize (T = 29.0 °C) | 0.10 | 12.79 | 0.38 | 1.39 | 1.28 | 8.69 | 0.63 | 1.62 | 18.49 | 14.59 |
| Maize ($\psi$ = 30 kPa) | 0.11 | 10.91 | 0.45 | 1.31 | 2.18 | 8.40 | 0.47 | 1.78 | 19.37 | 6.26 |
| Maize ($\psi$ = 200 kPa) | 0.11 | 8.62 | 0.53 | 1.01 | 1.72 | 6.84 | 0.45 | 2.09 | 21.75 | 5.66 |
| Maize ($\psi$ = 800 kPa) | 0.12 | 6.55 | 0.57 | 0.86 | 1.31 | 5.92 | 0.48 | 1.98 | 19.00 | 6.21 |
| Maize ($\psi$ = 1600 kPa) | 0.10 | 6.16 | 0.51 | 0.81 | 1.08 | 5.58 | 0.52 | 1.53 | 17.13 | 7.50 |
| Pea ($\sigma$ = 600 kPa) | 0.04 | 12.27 | 0.35 | 1.19 | 2.46 | 10.75 | 0.30 | 0.80 | 25.92 | 3.49 |
| Pea ($\sigma$ = 2000 kPa) | 0.06 | 7.86 | 0.23 | 1.83 | 1.36 | 9.98 | 0.72 | 0.12 | 4.447 | 19.26 |
| Tomato ($\sigma$ = 74 kPa) | 0.27 | 3.55 | 0.20 | 0.32 | 0.36 | 0.85 | 0.64 | 3.37 | 28.73 | 1.37 |
| Tomato ($\sigma$ = 194 kPa ) | 0.15 | 3.69 | 0.30 | 0.31 | 0.37 | 1.11 | 0.78 | 3.19 | 30.09 | 3.02 |
| Tomato ($\sigma$ = 323 kPa ) | 0.15 | 3.41 | 0.24 | 0.34 | 0.34 | 2.76 | 0.79 | 1.81 | 15.07 | 26.00 |
| Wheat ($\sigma$ = 74 kPa ) | 0.15 | 3.63 | 0.53 | 0.60 | 0.36 | 3.45 | 0.99 | 1.48 | 8.32 | 65.45 |
| Wheat ($\sigma$ = 194 kPa ) | 0.16 | 3.36 | 0.43 | 0.59 | 0.34 | 3.15 | 0.93 | 1.10 | 7.46 | 32.00 |
| Wheat ($\sigma$ = 323 kPa ) | 0.19 | 2.34 | 0.55 | 0.66 | 0.23 | 2.91 | 1.15 | 0.67 | 3.13 | 77.17 |

To compare how the different species differed in their growth strategies, we first plotted their normalised growth kinematics functions together. The normalisation of the growth velocity functions *g*(*x*) showed that Arabidopsis was the fastest growing of all species studied considering its small radius (Figure 5A). Its relatively large elongation zone was comparable to that of tomato, a similarly fast-growing species, but it also showed a large growth acceleration parameter *A*. By comparison, species such as pea or wheat exhibited slow root elongation rate relative to their size. Normalising distances by the length of the elongation zone collapsed all growth velocity functions onto a single curve, reducing interspecific variation to differences in the radius functions alone (Figure S2). This transformation revealed that species such as Arabidopsis and tomato taper much more rapidly, making them less capable of reducing friction than pea (Figure 5B). Simulations of roots rescaled to a 1 mm diameter further showed that tomato and wheat possess the tip shapes the least optimised for growth stability, unlike the root tips of Arabidopsis, which exhibited the smallest deflection.

**Figure 5:**
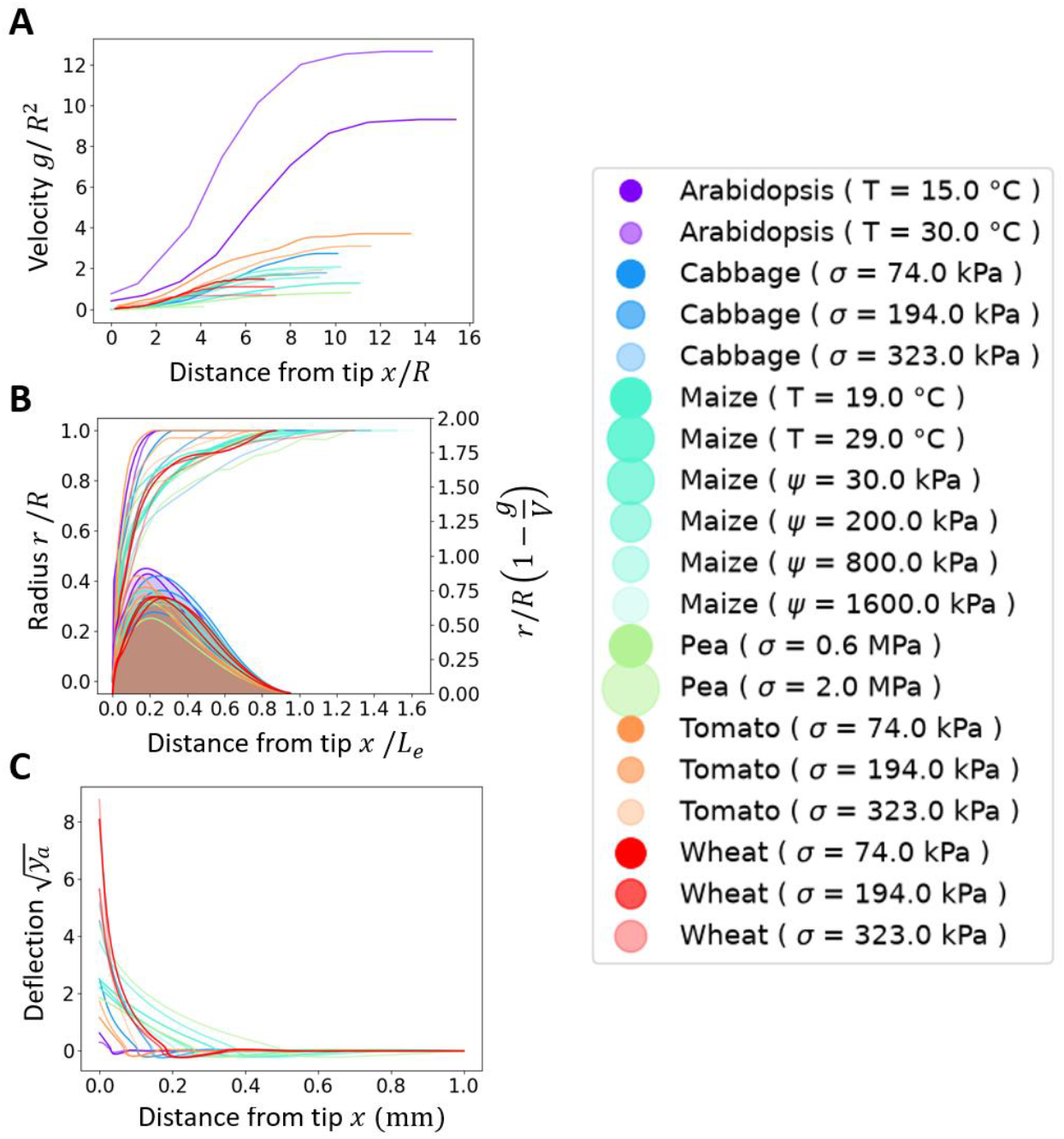
Morphology and growth kinematics across plant species and implications for the efficiency of growth. (A) The root growth velocity profile following normalisation by the root radius. Smaller root radii showed increased normalised velocity due to higher growth acceleration parameter *A* but less due to longer elongation zone *L*_*e*_/*R*. (B) When distances were normalised by the length of the elongation zone, pea and wheat plants displayed the sharpest tip profiles (non-shadowed curves, left y-axis) and experience the lowest friction, as shown in the friction attenuation factor linked to growth, 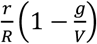 equation 13 (shadowed curves, right y-axis). (C) Simulations showed that, once the root diameter was normalised to 1 mm, Arabidopsis roots exhibited the smallest tip deflection which is consistent with the species experiencing the largest frictional resistance from the soil.

### Root growth strategy is determined by the root radius and soil mechanical properties

To understand how root growth strategies vary amongst species, we first looked at the emerging relationships between the multiplicative contrasts for velocity, friction, and stability of growth. Since sharper tips have a reduced second moment of inertia at the apex and are therefore more flexible, there was a strong correlation between the multiplicative contrasts for friction and stability (Figure 6A) but less so between velocity and friction or stability (Figure S3 and S4). A principal component analysis confirmed this trade-off, with the first principal component being dominated by the friction and stability contrast. Interestingly, this principal component also showed a strong correlation with radius (Figure 6A, inset), indicating that growth strategies centred on reducing friction were characteristic of larger roots, while strategies centred on stability were characteristic of smaller roots.

**Figure 6:**
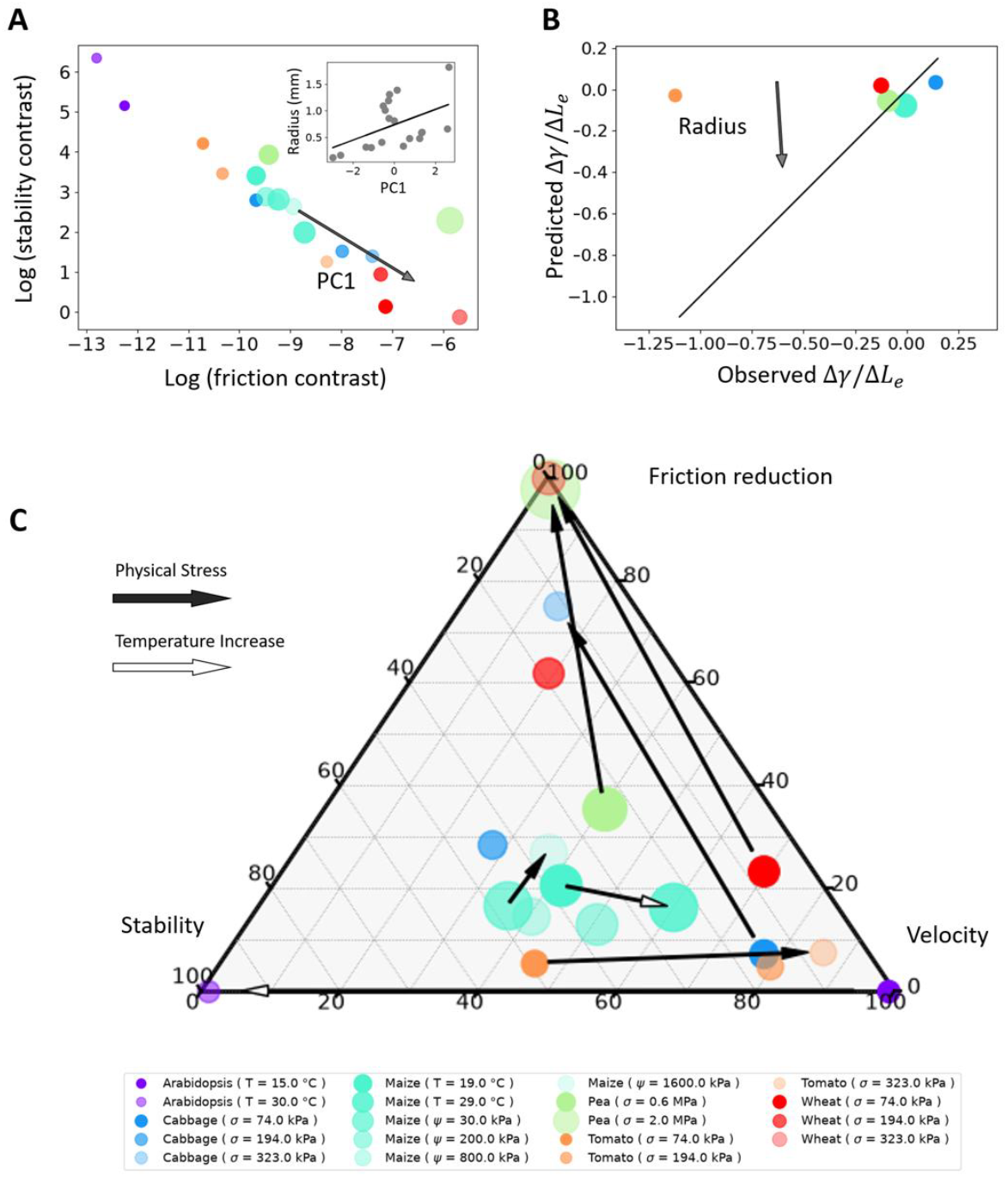
Growth strategies to overcome mechanical resistance from the soil. (A) The friction - stability trade-off gave the strongest indication of plants’ strategies to overcome mechanical resistance from the soil, as was indicated by a principal component analysis. The arrow indicates the first principal component (PC1). Plants with larger root radii favoured a reduction in the friction experienced during growth, but this came at the cost of reduced stability. Radius data was strongly correlated with PC1 (inset). (B) Plant responses to increased mechanical resistance from the substrate showed larger radius responded less strongly than other plants. (C) Overall, there were clear links between strategy for growth unconstrained and the response to the stress. Species such as tomato which tip shapes appeared less sensitive to deflection responded by safeguarding velocity (greater changes in tip sharpness than in the root elongation zone). In contrast, species such as maize which favoured velocity when unstressed responded by improving resistance to penetration. Changes to in responses to temperature did not appear correlated to improving the penetration resistance of growth.

Since data on root responses were obtained on varying conditions, we characterised the response of the root as the ratio between the changes in root tip sharpness (Δ*γ*) and the changes observed in the root elongation rate (Δ*L*_*e*_). Results showed that the experimentally observed responses matched model predictions qualitatively. The root tip tended to sharpen (Δ*γ* > 0) and the length of the elongation zone reduced (Δ*L*_*e*_ < 0). Quantitatively however, the model did not predict well the responses of the root (Figure 6B), with large deviations observed notably for tomato roots which exhibited a much stronger sharpening of the root tip than expected. Observed adjustments made to the tip shape Δ*γ*, in the range of −0.07 to 0.4 were larger than predicted (from −0.01 to 0.02). Similarly, the observed adjustments made to the root elongation zone Δ*L*_*e*_, in the range −4.7 mm to −0.1 mm, were larger than predicted (from −0.3 mm to −0.2 mm). It is interesting to note here that deviations from expected responses were related to the root radius. Roots with larger radii exhibited smaller adjustments in root-tip shape compared with species that had thinner roots.

Finally, we collapsed all the root responses data into a ternary plot with velocity, stability and friction contrasts as apices (Figure 6C). The plot revealed a general trend in which plant roots respond to mechanical resistance from the soil by reducing the frictional resistance experienced at the root tip, either by reducing the elongation rate or by reducing tip sharpness. Species such as pea or tomato, whose growth strategies emphasise growth velocity, tend to compensate by adjusting stability and friction. Species whose strategies prioritize growth stability respond by rebalancing velocity and friction instead. Water stress, which closely intertwin with mechanical stress, triggered the same pattern of response for maize roots. By contrast, increased temperature did not appear related to reducing resistance to penetration.

## Discussion

### Root tips and soil physical constrains to growth

When roots grow into soil, they must deform the surrounding soil particles to create the space they occupy. Achieving this requires plant roots to produce the energy necessary to overcome friction and the compressive forces acting on the outer tissues. Friction is the most notable of these two forces. Not because friction energy systematically exceeds the energy lost to deform the soil radially, but because it is the factor a root can most easily act upon. This is captured in equations 12 and 13 which show the energy required to compress the soil is independent on root growth kinematics. The model does not incorporate a dependence of the mechanical energy of soil deformation on growth rate, and it is possible that slow growth may reduce compression forces. Nevertheless the myriad of mechanisms which roots employ to reduce friction, including the production of border cells, release of mucilage (Bengough *et al*. 2006) or circumnutations (Leuther *et al*. 2025) gives weight to that hypothesis.

A second and perhaps less understood physical constraint to root growth is soil heterogeneity. Heterogeneity can arise from numerous sources, including variation in soil texture, differences in soil structure, aggregation properties, or the presence of cracks and pores (Jin *et al*. 2013). As roots grow through soil, they must navigate this spatially disorganised environment, encountering zones of contrasting resistance. Plant roots can benefit from heterogeneity, for example by growing through zones of reduced resistance (Landl *et al*. 2019). But soil heterogeneity also forces roots to alter their growth direction, reducing the efficiency with which they proliferate vertically by increasing the tortuosity of the root path (Dupuy *et al*. 2018).

What this study indicates, is that the shape of root tips reflects the necessity for plants to overcome both physical constrains. When soil mechanical resistance increases, the resulting forces acting on the root slow down cell expansion (Dexter 1987). Sharper root tips can partially mitigate this effect by reducing frictional resistance as the root elongates. The effects of friction at the root tip were observed from experiments on decapped root tips and mutant plants (M Iijima *et al*. 2003; Morio Iijima *et al*. 2003; Vollsnes *et al*. 2010). Using penetrometer tests with rotating probes, it was also possible to isolate the frictional component of penetration resistance and gain insight into how plant roots may circumvent this constraint (Mckenzie *et al*. 2013; Leuther *et al*. 2025). An analysis of genotypic variation in wheat root tip shape showed that tip sharpness correlates positively with elongation rates in compacted soils (Colombi *et al*. 2017).

It is also well established that plant roots are prone to buckling and bending when they encounter significant axial resistance from the soil (Whiteley *et al*. 1982; Silverberg *et al*. 2012; Bizet *et al*. 2016). The increased confinement of the soil restricts lateral soil displacement, leading to bending events with higher curvature (Martins *et al*. 2019), and for this reason soil compaction is also associated with an increase in root tortuosity (Popova *et al*. 2016; Correa *et al*. 2019). Root stiffness is therefore a critical factor in preventing tortuosity, and the root tip is the most relevant region in this respect, as it is where the radius tapers.

### Smaller roots prioritise stability and speed, larger roots prioritise friction

Plant species show substantial variation in the form and development of their root tips (Ganesh *et al*. 2022), and it is insightful to consider how these differences influence a plant’s ability to overcome physical constraints imposed by the soil. The analysis of the growth kinematics parameters of the roots of various species revealed that the radius of the root was a main driver for changes in growth strategy (Figure 6A). Plant species such as Arabidospsis, tomato or cabbage, which had smaller root radii, exhibited features that favoured stability. They showed a rapid tapering the root radius at the tip (Figure 5B) which induced more friction but made the root more resistant to bending (Figure 5C).

This feature was also associated with a larger relative growth zone, making them fast considering their small radii (Figure 5A). Species with larger root radii on the other hand, had a larger proportion of their elongation zone present in the tapering part of the root tip, making them more efficient mechanically, but also less stable relatively to their radius. Larger diameter may experience increased compression and friction stresses than predicted by the model, making larger plants more susceptible to friction loss. These results also go against the study from He et al. (2022), which found a positive relationship between root tip angle and root radius. To resolve this apparent contradiction, greater emphasis must be placed on standardising measurements of root tip geometry, particularly by accounting for the length of the growth zone when fitting radius functions.

All plants showed a reduction in the growth zone and often resulted in root tips growing sharper, in responses consistent with a reduction of friction. This pattern was previously reported by Kirby and Bengough (2002) and Colombi et al. (2017), and our results consolidated their observation across a broader range of species. We also observed the well documented increase in root radius (Potocka and Szymanowska-Pułka 2018), and found additionally that root radius was a principal factor explaining plant responses to mechanical stress. Plants with larger radii responded less strongly to the mechanical stress than plants with smaller radii (Figure 6B). Overall, the responses observed reflected the plant’s effort to maintain the root elongation rate, suggesting that primary roots may prioritise exploratory functions over exploitative ones when under mechanical stress.

The nature of soil heterogeneities, which are not scale-invariant (Yudina and Kuzyakov 2023), may explain the observed overall gradient in root developmental strategy. Smaller roots may be more capable of finding small pores to grow unimpeded, which is consistent with a fast and high friction growth strategy. Larger roots, however, may be less likely to find path of least resistance through the soil and may be more exposed to friction. A reduced radius also makes roots more sensitive to deflection, as root rigidity scales with the fourth power of radius (Kolb *et al*. 2017), whereas traits such as elongation rate and surface area scale linearly with radius, and biomass scales with the square root of the radius. Blunt root tips may prevent excessive bending and deflections when penetrating soil from a pore, which confirms observation from Roue et al. (2020).

### The economics spectrum of root establishment

Root growth strategies have been characterised based on tissue density, chemical composition and anatomical features, following the now classic Root Economics Spectrum (Prieto *et al*. 2015; Roumet *et al*. 2016). At one hand of the spectrum, roots with high specific root length, small radii, and low tissue density are cheaper to construct and allow fast root proliferation. These species are adopting an acquisitive strategy. In contrast, longevity is associated with slow-growing roots that are denser and larger in radius, reflecting a conservative resource-use strategy. A number of studies have both confirmed or contradicted the Root Economics Spectrum (Kramer-Walter *et al*. 2016; McCormack and Iversen 2019) and others have sought to expand the dimensionality of the trait space, for example by adding a collaboration gradient that reflects a root’s capacity to form associations with fungi (Bergmann *et al*. 2020).

Because root growth is physically constrained by the substrate, many of the traits traditionally associated to fast growth may not directly relate to an ability to overcome the mechanical impedance from the soil. Low-density tissues may notably lack the stiffness required to resist buckling which would in turn prevent soil penetration (Kolb *et al*. 2017; Vanhees *et al*. 2020). Additionally, many of the root traits commonly assessed within the Root Economics Spectrum change as roots mature. Secondary growth, starch accumulation or programmed cell death of the cortex for example, root radius, tissue density and specific root length, change during the lifespan of the plant (Drew *et al*. 2000; McCormick *et al*. 2013; Lyu *et al*. 2025). Cell wall thickening profoundly alters the chemical composition of root tissues, adding various types of lignin, cellulose, pectin and hemicellulose to the apoplast (Cantó-Pastor *et al*. 2025). As a result, traits that allow fast expansion in soil may not be those measured in field experimentation which usually target mature roots.

To better understand root growth strategies, it is therefore useful to frame developmental traits within an exploration–exploitation paradigm (Camenzind *et al*. 2024). Since the elongation rate is largely controlled by the radius of the root (Cahn *et al*. 1989; Pagès 1995), larger roots are therefore leaning on the explorative end of the spectrum. Thinner roots, which expand more slowly into the soil but can produce a greater number of roots, are more capable of exploiting resources intensively and are therefore on the exploitative end of the spectrum (Figure 7). The energetics of growth is independent of the exploration-exploitation axis since it is not dependent on root radius but on the regulation of axial and radial cell expansion in the root apical meristem. Sharper root tips are more capable of reducing the energy lost to friction, but this comes at the cost of an increased likelihood of deflection in heterogeneous substrates. Consequently, trade-offs along the biomechanical axis emerge from the interplay between frictional losses and inefficiencies linked to the stochasticity of growth.

**Figure 7:**
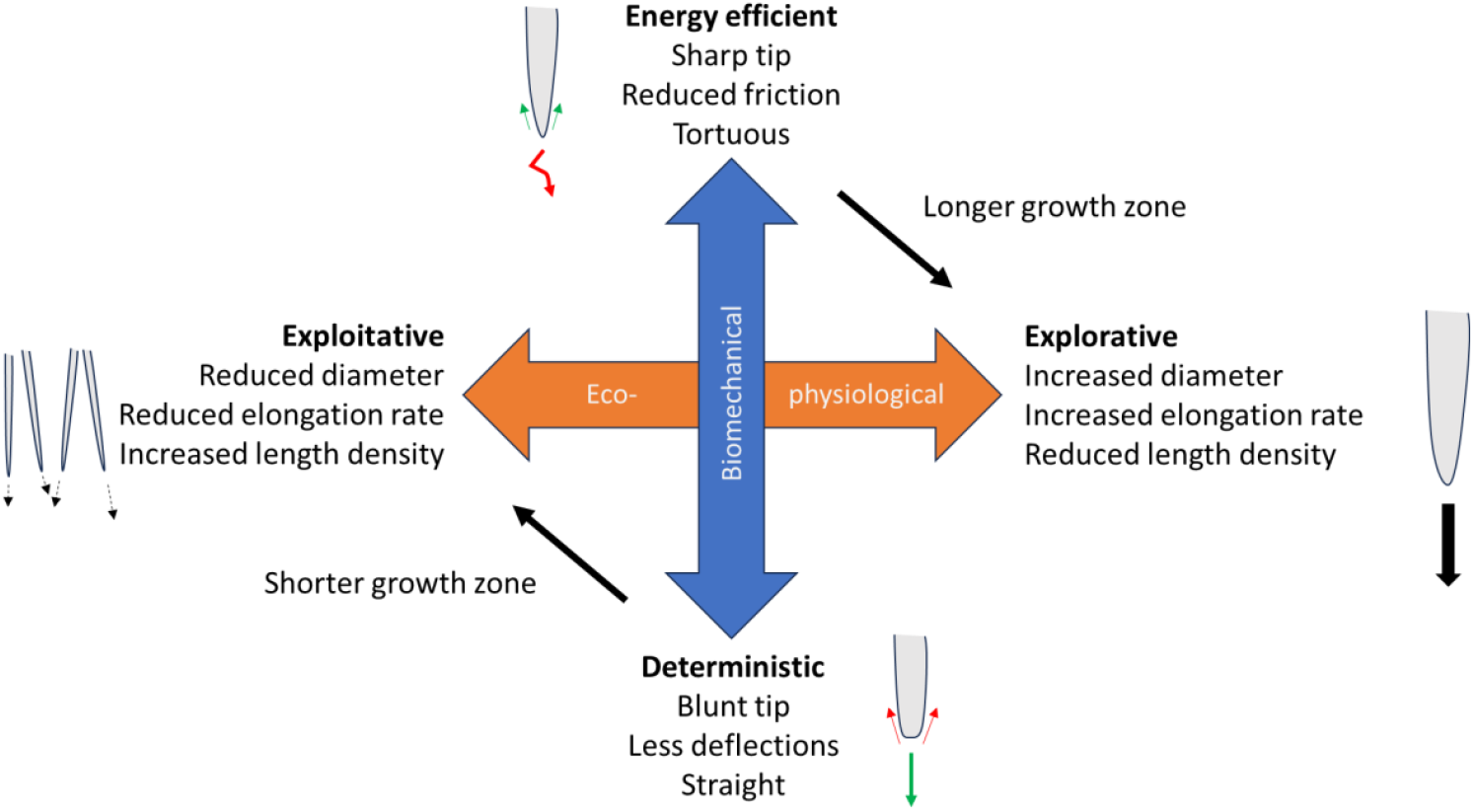
Root growth strategies in a mechanically impeding medium. Amongst root systems of equal biomass and radius, two distinct axes characterise growth strategies. Root systems which maintain high root elongation rate can do so by increasing the root radius for a more explorative strategy (orange arrow, right). This is at the expense of the number and length density of roots, which can be maintained by keeping the root radius unchanged or even reduced (orange arrow, left). The later strategy is exploitative. Root growth kinematics functions (here the function *r* and *g*) are indicative of the mechanical efficiency of growth (blue arrow). Sharper root tips (top) reduce energy loss due to friction, but this advantage comes at the cost of reduced mechanical stability. Such roots are prone to frequent deflections and tortuous growth trajectories. Adjustment to the root elongation zone affects the energy loss due to friction but also allows adjustments along the exploitative-explorative axis (oblique arrows).

## List of symbols

| Symbol | Description | Value |
| --- | --- | --- |
| $x$ | Distance from the root tip | mm |
| $\theta(x)$ | Angle of the root envelope | |
| $r(x)$ | Radius function | mm |
| $R$ | Root radius | mm |
| $V$ | Root tip velocity (also root elongation rate) | mm h<sup>-1</sup> |
| $g(x)$ | Growth velocity function | mm h<sup>-1</sup> |
| $\partial_x g(x)$ | Strain rate | h<sup>-1</sup> |
| $\mathbf{v}(x)$ | Growth velocity vector | mm h<sup>-1</sup> |
| $\mathbf{V}(x)$ | Global root velocity vector | mm h<sup>-1</sup> |
| $\bar{\mathbf{V}}(x)$ | Root-soil slip velocity vector | mm h<sup>-1</sup> |
| $\mathbf{e}_x, \mathbf{e}_y$ | Horizontal and vertical vectors | |
| $\mathbf{n}, \mathbf{t}$ | Normal and tangential vectors of the root envelope | |
| $\sigma_n$ | Soil yield stress | Pa |
| $\tau$ | Component of the penetration resistance attributed to friction | Pa |
| $\mu$ | Friction coefficient | |
| $W$ | Mechanical power needed to form a root cavity | W |
| $F$ | Force exerted by the root on the soil | N |
| $'$ | Non dimensional variable | |
| $L_e$ | Length of the root growth zone | mm |
| $A$ | Growth acceleration parameter | mm<sup>-1</sup> h<sup>-1</sup> |
| $\alpha$ | Growth acceleration zone parameter | |
| $\gamma$ | Tip sharpness parameter | |
| $L_r$ | Length of the root tapering zone | mm |
| $K$ | half-saturation constant for the radius function | mm<sup>γ</sup> |
| $E_\beta$ | Cost function | |
| $\beta_v, \beta_f, \beta_\theta$ | Weights associated with penalties for reduced velocity, increase friction and stability trade-offs | |

## Supplemental Files

Supplementary Documents.pdf

## Data availability

The dataset generated in this study is available at https://doi.org/10.5281/zenodo.19697131

## Artificial Intelligence Generated Content

No AI system was used for data analysis, interpretation of results, generation of scientific conclusions or to generate text. Language editing tools have been used for proofreading the manuscript.

## Author contributions

Conceptualization: LXD

Methodology: LXD

Investigation: LXD, JY, GH

Funding acquisition: LXD

Project administration: LXD

Supervision: LXD

Writing – original draft: LXD

Writing – review & editing: LXD, JY

