## Supplementary Information for "Growth under constraints: root tip development controls trade-offs between speed and mechanical efficiency"

**Supplementary Method**

*The optimal root growth kinematics*

Root growth kinematics parameters $L_e$ and $\gamma$, noted as vector $\boldsymbol{\alpha} = (L_e, \gamma)$, determine the biomechanical efficiency of growth in three ways: the growth velocity $V = g(L)$, the energy lost by friction $\sigma_f$, and the ability of the root to grow straight, represented by the root tip deflection angle $y_a$ in response to a force, here chosen arbitrarily as 5 mN. We assume growth and morphology of the root tip result from a trade-off among the penalties imposed by each of these processes. The overall cost associated with a given growth strategy is therefore defined as an error function $E_{\boldsymbol{\beta}}(\boldsymbol{\alpha})$ which we assume to be the sum of the penalty for slow growth, the penalty for the resistance to friction and the penalty for deflection of the root tip, $E_{\boldsymbol{\beta}} = \beta_v V^{-2} + \beta_f \tau^2 + \beta_\theta y_a^2$, which for simplicity we write as $E_{\boldsymbol{\beta}} = \sum \beta_i E_i$. $\beta_i, i < m$ are weighting factors termed penalty coefficients. The optimal root tip parameters $\boldsymbol{\alpha}^* = (L_e^*, \gamma^*)$ are those that minimizes the error function, such that

$$\nabla E_{\boldsymbol{\beta}}(\boldsymbol{\alpha}^*) = \sum \beta_i \nabla E_i(\boldsymbol{\alpha}^*) = \mathbf{0} \qquad (1)$$

In matrix form, this can be rewritten as $[\mathbf{G}]\boldsymbol{\beta} = \mathbf{0}$. $\boldsymbol{\beta}$ indicates a column vector of weights in the error function, and $[\mathbf{G}] = \left[\partial_i E_j\right]_{i \le n, j \le m}$ is the gradient matrix. Since $m > n$, we determine the first $n$ weights while fixing all the others to 1,

$$\boldsymbol{\beta}_{\mathbf{1,n}} = -\left[\mathbf{G}_{\mathbf{1,n}}\right]^{-\mathbf{1}}\left[\mathbf{G}_{\mathbf{n,m}}\right]\mathbf{I}, \qquad (2)$$

with

$$\boldsymbol{\beta}_{\mathbf{k,l}} = [\beta_i]_{k \le i \le l} \quad \left[\mathbf{G}_{\mathbf{k,l}}\right] = \left[G_{i,j}\right]_{i \le n, k \le j \le l}$$

To determine how a root should adapt to maintain the same growth strategy when soil conditions are changing, we determined how the growth kinematic parameters vary in order to remain on the minimum of the error function. If soil parameters $\sigma_n$ and $\mu$ change from reference values $s_0$ to increased values $s_1$, the optimal root tip parameters noted $\boldsymbol{\alpha}^*$ must change parameters $\boldsymbol{\alpha}^{**}$ to keep minimising $E_{\boldsymbol{\beta}}$. The Taylor approximation of $E_{\boldsymbol{\beta}}$ is therefore as follow

$$E_{\boldsymbol{\beta}}(\boldsymbol{\alpha}^{**}, s_1) = \mathrm{E}_{\boldsymbol{\beta}}(\boldsymbol{\alpha}^*, s_1) + \mathbf{D}(\boldsymbol{\alpha}^{**} - \boldsymbol{\alpha}^*) + \frac{1}{2}(\boldsymbol{\alpha}^{**} - \boldsymbol{\alpha}^*)^{\mathrm{T}}[\mathbf{H}](\boldsymbol{\alpha}^{**} - \boldsymbol{\alpha}^*). \qquad (3)$$

$\mathbf{D}$ and $[\mathbf{H}]$ are the gradient vector and Hessian matrix for the error function $\mathbf{E}$ evaluated with regards to the growth kinematic parameters for soil conditions $s_1$. For the new set of root tip parameters $\boldsymbol{\alpha}^{**}$ to be optimal with error function $\mathbf{E}$ the gradient of the error function must equal the null vector,

$$\nabla E_{\boldsymbol{\beta}}(\boldsymbol{\alpha}^{**}) = \mathbf{D} + \frac{1}{2}[\boldsymbol{A}](\boldsymbol{\alpha}^{**} - \boldsymbol{\alpha}^{*}) = \mathbf{0} \tag{4}$$

Where $[\mathbf{A}] = [\mathbf{H}]^{\boldsymbol{T}} + [\mathbf{H}]$.

Therefore, if the strategy of the plant remains unchanged, root tip traits would vary to follow the condition

$$\boldsymbol{\alpha}^{**} - \boldsymbol{\alpha}^{*} = -2[\mathbf{A}]^{-\mathbf{1}}\mathbf{D}. \tag{5}$$

**Supplementary Figures**

Figure S1. Root growth kinematics analysis. (A) High-throughput live imaging system was used to capture time-lapse images of roots grown using agarose gels at concentrations 1.5%, 3.0%, and 4.5%. (B) Raw images (left) were processed using KineRoot software to quantify root elongation rate and diameter (right). Elongation was determined by tracking manually annotated reference point (red) over time, while diameter was determined from the root contour (green and yellow lines). The scale bar is 0.2 m. (C–D) Resulting elongation rate and diameter were quantified as functions of distance from the root tip.

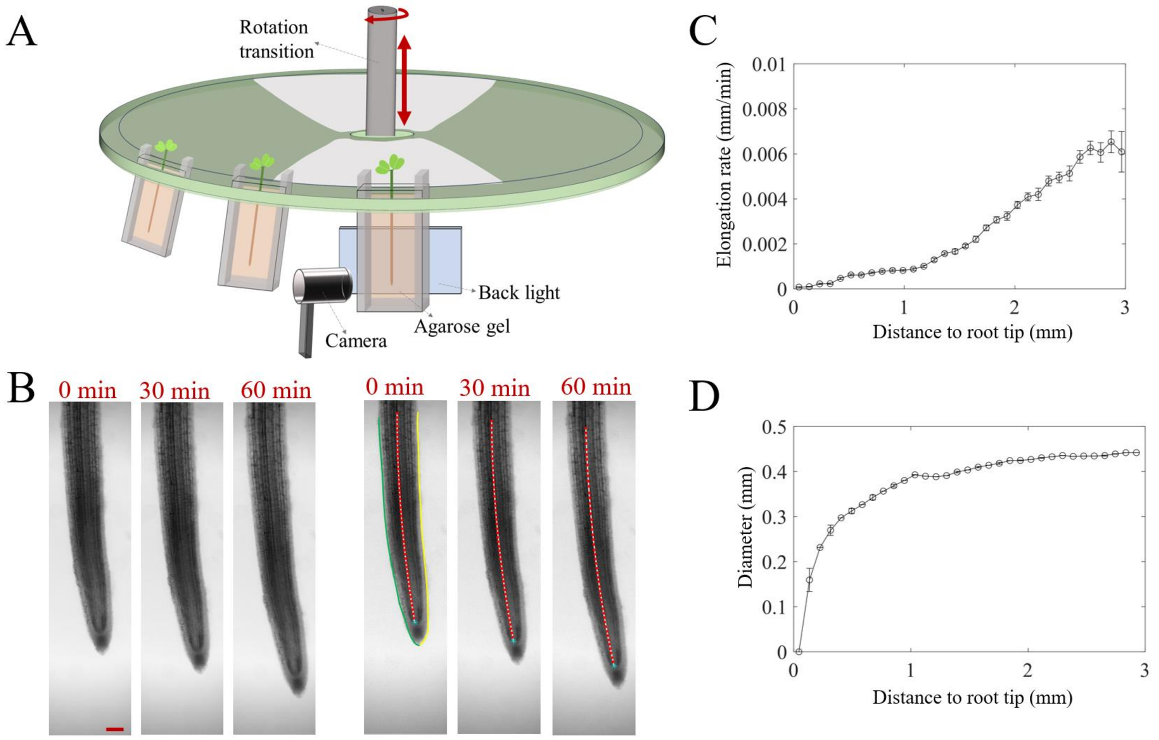

Figure S2. Cells' axial velocity after normalisation.

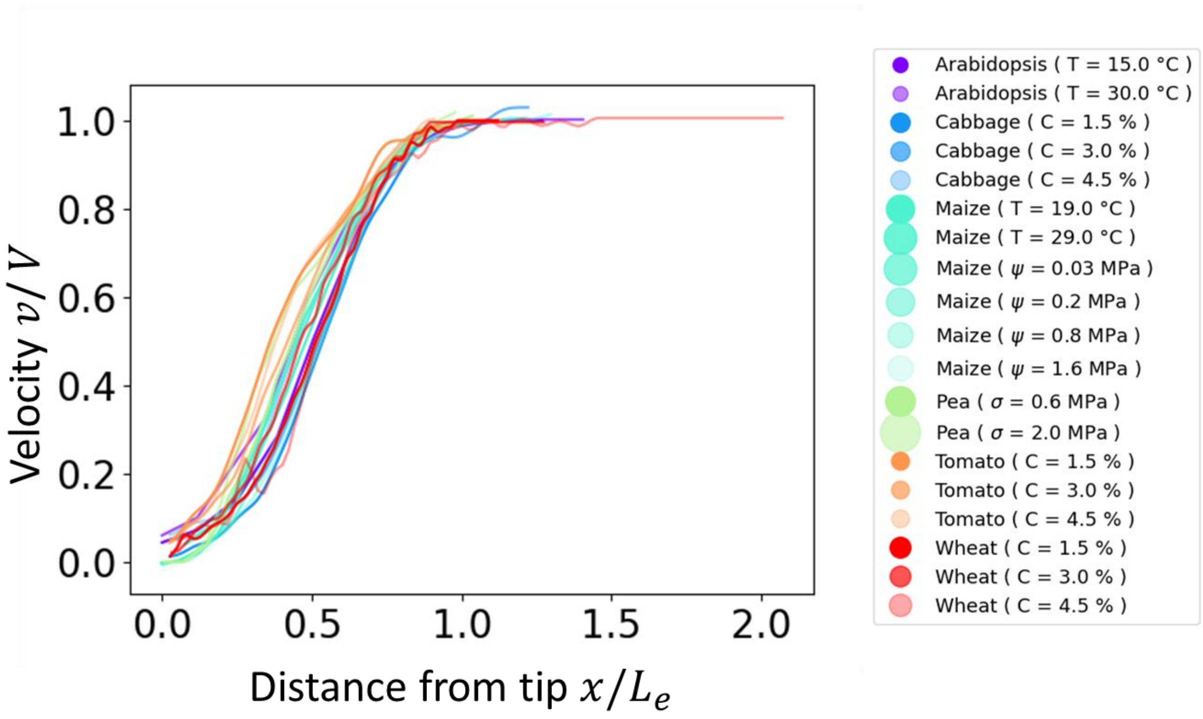

Figure S3. Relationships between the multiplicative contrast for velocity and stability.

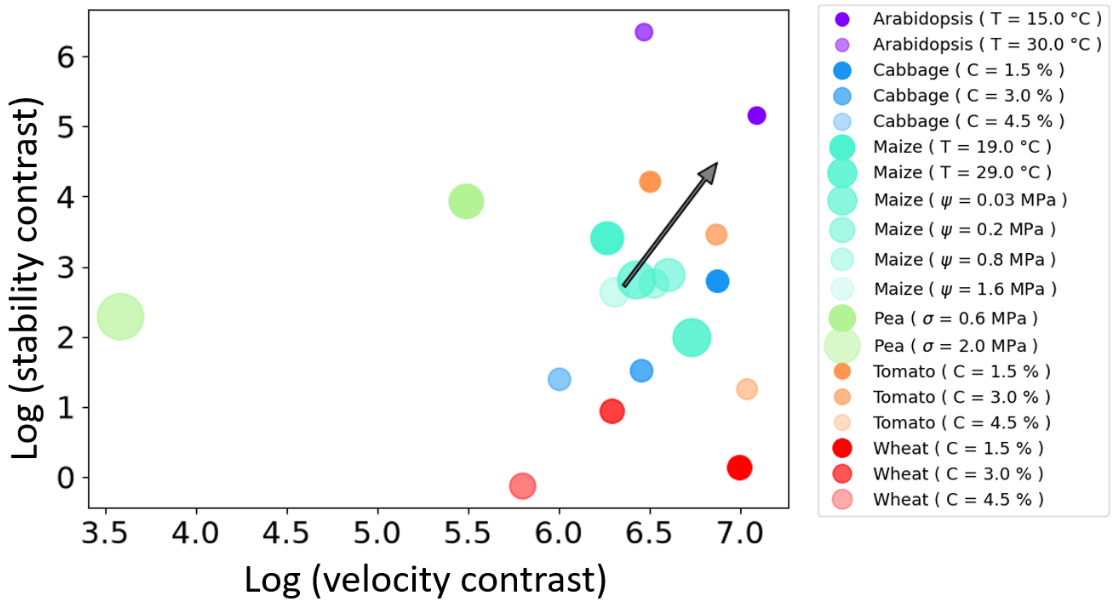

Figure S4. Relationships between the multiplicative contrast for velocity and friction.

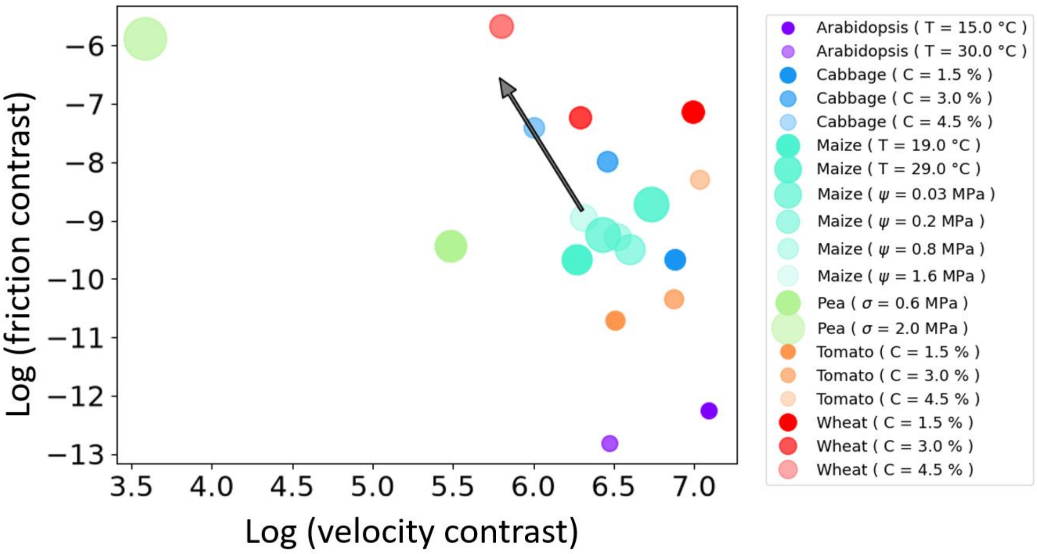
